# Extra-sheath transplantation of human retinal organoids (hROs): effects on visual recovery and retinal repair following optic nerve injury

**DOI:** 10.64898/2026.11.18.767615

**Authors:** Yushan Luo, Xia Chen, Huan Qin, Jiahui Kang, Kai Yao, Haiwei Xu

## Abstract

Traumatic optic nerve injury interrupts retinal ganglion cell output and can produce permanent visual disability, yet a therapeutic strategy that couples retinal protection with recovery of visual-pathway activity remains unavailable. Here, we evaluated human retinal organoid transplantation at the site of optic nerve crush in C57 mice. Human retinal organoids derived from embryonic stem cells were used at day 30 and showed retinal-lineage immunoreactivity together with spontaneous microelectrode-array activity. Across post-surgical days 7, 14, and 28, transplantation was associated with preservation of optic-disc appearance and ganglion cell layer thickness. The structural benefit became statistically distinguishable from the crush group at day 28 and remained incomplete relative to the uninjured or sham group. Serial RBPMS and TUJ1 immunostaining further supported attenuation of retinal ganglion cell body and neurite loss, while IBA1 staining indicated an altered microglial response. Full-field electroretinographic b-wave responses showed no detectable between-group differences, arguing against a major outer-retinal effect. In contrast, flash visual evoked potential P2 amplitude was higher in transplanted than in crush-only mice at all measured time points, consistent with improved visual-pathway responsiveness. Transplant recipients also showed enhanced optomotor responses across tested spatial-frequency and contrast conditions. Light-dark behavior shifted toward greater dark-compartment occupancy and activity, although this assay remains non-specific for visual recovery. Together, these findings support a model in which hROs transplantation at an optic nerve injury site promotes multi-level functional preservation after optic nerve crush. Definitive evidence for donor-derived circuit reconstruction will require graft tracing, synaptic validation, and prespecified electrophysiologic analyses.

**Graphical abstract:** 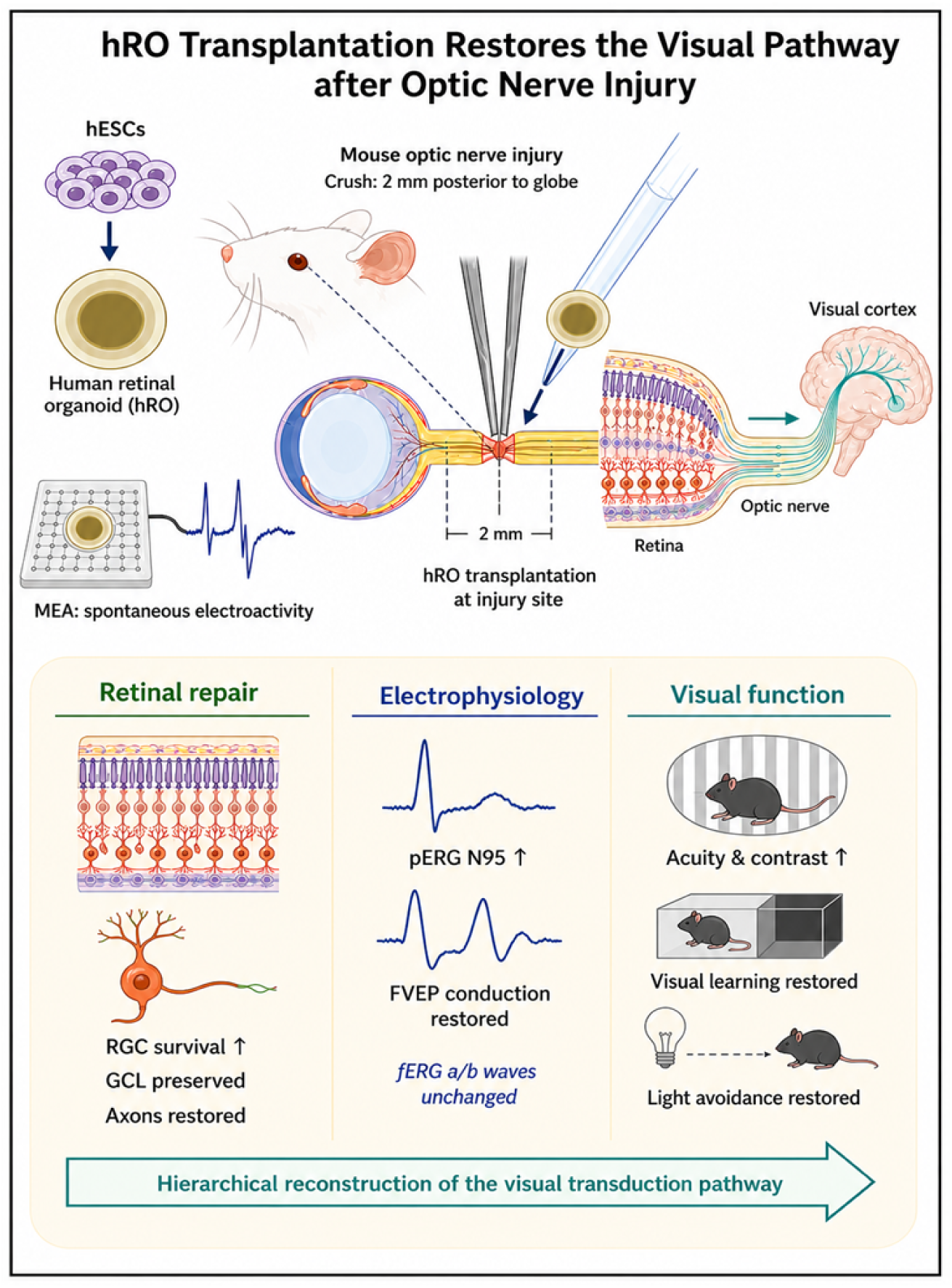

## Introduction

Traumatic optic neuropathy (TON) disrupts retinal ganglion cell (RGC) axons and can cause permanent visual disability. The primary axonal insult is followed by delayed loss of RGC somata and neurites within the retina, together with inflammatory and glial responses that can amplify secondary injury^1, 2^. The adult optic nerve remains a powerful model for central nervous system repair. PTEN deletion, combined PTEN and SOCS3 manipulation, and approaches that improve conduction in regenerated axons have each advanced the field^3–6^. These studies also reveal a critical distinction: anatomical outgrowth, RGC survival, and physiologically useful visual transmission are related but non-equivalent outcomes^6–8^. The injured retina is therefore not merely the origin of damaged axons; it is a therapeutically accessible compartment in which preserving surviving RGCs and stabilizing the local microenvironment may retain residual visual-pathway function.

Current experimental strategies increasingly target this retinal-protection window. Molecular and pharmacological interventions can enhance RGC survival by modifying intrinsic stress responses, trophic signaling, oxidative stress, or inflammatory signaling. Biomaterial and nanoparticle platforms further enable local, sustained delivery of neuroprotective cargo. For example, brimonidine-loaded albumin nanoparticles improved RGC survival after optic nerve crush, and polydopamine nanoparticles reduced retinal oxidative stress and microglial activation while attenuating RGC loss^9, 10^. These studies establish the value of local retinal neuroprotection, but single-agent approaches are constrained by finite payloads and usually address only one component of the post-injury microenvironment. Cell-based interventions offer another complementary strategy because they can provide sustained trophic support and modulate the injury milieu. Mesenchymal stromal-cell transplantation and human periodontal ligament-derived stem-cell transplantation improved host RGC survival after optic nerve crush in preclinical models, with evidence implicating secreted trophic factors and local cell-cell interactions^11, 12^. However, cell therapy that preserves host RGCs should be distinguished from claims of donor-derived retinal replacement or reconstructed eye-to-brain connectivity. Direct evidence of donor survival, axon extension, synapses, and target innervation is required before invoking the latter mechanism.

Human retinal organoids offer an alternative therapeutic resource because they can self-organize neural-retinal tissue from pluripotent stem cells and reproduce important features of retinal lineage specification, architecture, and functional maturation^13–17^. Here, we used human retinal organoids (hROs) derived from human embryonic stem cells and transplanted them at the optic nerve crush (ONC) site in C57 mice. The study integrates donor-tissue quality control, serial in vivo retinal imaging, RGC and glial immunohistology, full-field and pattern electroretinography, flash visual evoked potentials, and visually guided behavioral assays. OCT and electroretinographic changes after mouse ONC have been characterized previously^18, 19^, while pattern ERG and VEP provide complementary readouts of inner-retinal and post-retinal visual function^20–22^. Optomotor testing offers a sensitive measure of spatial vision in mice^23^. In contrast, the light-dark box measures light-context exploration and aversion rather than visual acuity perse^24^. Our primary interpretation is that extra-sheath hROs transplantation may preserve the injured host retina and improve residual visual-pathway responsiveness

## Materials and Methods

### Animals and experimental design

Work involving human embryonic stem cell-derived retinal organoids was approved by the Institutional Animal Care and Use Committee of Army Medical University and complied with the ARVO Statement for the Use of Animals in Ophthalmic and Vision Research and relevant institutional guidelines.

Animals were allocated to sham, optic nerve crush (ONC), and ONC with extra-sheath human retinal organoid transplantation (Trans) groups. Fundus photography, optical coherence tomography (OCT), electrophysiological recording, and behavioral testing were performed at the indicated post-operative time points. Retinal tissue was collected on post-operative days 7, 14, and 28 for immunofluorescence analysis. Group allocation was randomized, and investigators responsible for imaging quantification, electrophysiology, and behavioral scoring were blinded to treatment allocation.

### Generation and characterization of human retinal organoids

Human retinal organoids (hROs) were generated from human embryonic stem cells (hESC) using a stepwise three-dimensional retinal differentiation protocol. Human retinal organoids for transplantation were generated using a classic SFEBq protocol (serum-free embryoid body culture with quick aggregation). Human embryonic stem cells were differentiated through neural induction and retinal specification in BMP4. Emerging retinal organoids were transferred to low-attachment culture plates and maintained in retinal maturation medium until transplantation.

Day-30 hROs were selected for transplantation on the basis of retinal morphology and structural integrity. Donor organoids were characterized by immunofluorescence staining for RAX and ISL1. Before transplantation, spontaneous electrophysiological activity of hROs was assessed by multielectrode-array recording using 3Brain BrainWave. Individual organoids were recorded for 3 minutes, and spontaneous extracellular activity was analyzed using spike-detection>8SD. Organoids or organoid fragments measuring 0.4mm^3^ were maintained until transplantation.

### Optic nerve crush and hROs transplantation

Mice were anesthetized with an intraperitoneal injection of 1.25% tribromoethanol (200 mg/kg; Avertin). After topical application of 2% Lidocaine, a small conjunctival incision was made in the left eye. Orbital soft tissue was gently separated to expose the intraorbital optic nerve while preserving the retinal blood supply. The optic nerve was crushed 2 mm posterior to the globe using Sterile tweezers for 30s.。

In ONC+hRO mice, one hRO graft (0.4mm) was placed at the exposed optic nerve lesion site using Syringe. The exact graft position was external to the optic nerve sheath. The conjunctiva was repositioned, and topical cyclosporine A (Neoral, Novartis, Basel) was applied. Sham-operated mice underwent optic nerve exposure without nerve crush. To prevent graft rejection, cyclosporine A (Neoral, Novartis, Basel) was administered intraperitoneally at a dose of 5 mg/kg, starting 1 day before surgery until 3 months after surgery.

### Fundus photography and OCT

Fundus photography and spectral-domain OCT were performed on post-operative days 7, 14, and 28. Mice were anesthetized, pupils were dilated with tropicamide eye drops (Santen), and corneal hydration was maintained with PBS. Color fundus photographs were acquired using Heidelberg Spectralis (Heidelberg Engineering) under identical acquisition settings for all groups. Fundus images were used for qualitative assessment of optic-disc border definition, optic-disc color, and retinal vascular appearance.

OCT images were acquired using the Spectralis OCT system (Heidelberg Engineering, Germany). B-scans were centered on the optic nerve head, and ganglion cell layer (GCL) thickness was measured at the optic disc region using manual segmentation method. For each eye, three scans were averaged to obtain one value. Image acquisition and GCL measurement were performed by investigators blinded to group allocation.

### Retinal immunofluorescence and image analysis

At post-operative days 7, 14, and 28, mice were euthanized using cervical dislocation, and eyes were enucleated. Retinal tissues were fixed in 4% paraformaldehyde for 1h and processed as retinal flat mounts or 15-μm cryosections. Samples were permeabilized and blocked in [buffer composition] and incubated with primary antibodies against RBPMS, β III-tubulin (TUJ1), IBA1, CD68, and GFAP. After washing, samples were incubated with species-appropriate fluorescent secondary antibodies and counterstained with DAPI.

Images were acquired using a Zeiss LSM 880 confocal microscope with identical laser power, detector gain, and exposure settings across experimental groups. RBPMS-positive cells were quantified as retinal ganglion cell-associated somata. TUJ1-positive neurite length and morphology were quantified using ImageJ and graphpad. IBA1-positive area, CD68-positive signal, and GFAP-positive area were measured using a prespecified thresholding workflow in ImageJ. For each retina, [number] fields from [central, mid-peripheral, and peripheral] regions were analyzed and averaged to generate one biological replicate.

### Electroretinography and flash visual evoked potentials

Full-field electroretinography (fERG) and flash visual evoked potentials (FVEPs) were recorded at 7, 14, and 28 days postoperatively. Mice were anesthetized and placed on a temperature-controlled platform. For fERG, mice were dark-adapted for 8h, pupils were dilated, and corneal electrodes were positioned on the tested eye. Reference and ground electrodes were placed at the forehead and tail, respectively. Full-field flashes were delivered using MAYO ERG, Japan and a-wave and b-wave amplitudes were measured using predefined waveform criteria.

For FVEP recording, the active electrode was positioned over the primary visual cortex at the stereotaxic coordinates or anatomical landmark used in the preceding TON transplantation study, with reference and ground electrodes at tail. Full-field flashes of the same intensity and frequency, with the same number of sweeps, as those used in the preceding TON transplantation study were delivered to the tested eye. Signals were amplified, filtered, digitized, and averaged. N1 latency, P2 latency, and N1–P2 peak-to-peak amplitude were quantified by an investigator blinded to group allocation.

### Behavioral assessments

Visually guided behaviors were assessed 2 weeks postoperatively using optomotor-response (OMR) testing, the Light–Dark Box test, a water-maze-based visual discrimination task, and a light-avoidance shuttle-box task. All behavioral tests were performed in a quiet testing room under standardized environmental conditions. Animals were allowed to acclimate to the testing room for 24 h before behavioral assessment.

For OMR testing, unrestrained mice were placed on a central platform surrounded by computer monitors displaying rotating vertical sine-wave gratings. Direction-selective head-tracking responses to clockwise and counterclockwise gratings were recorded using camera and tracking software. Spatial visual acuity was determined by increasing spatial frequency at 100% contrast until tracking responses were no longer detected. Contrast sensitivity was determined by reducing grating contrast at a fixed spatial frequency of 0.064 cycles/degree (cpd). The mean response threshold from both rotation directions was used for analysis.

Light-context behavior was assessed in a two-compartment Light–Dark Box consisting of illuminated and dark chambers connected by a doorway. Light intensity in the illuminated chamber was set to 300, 900, or 1,500 lux. Each mouse was placed in the starting compartment and allowed to explore the apparatus for 10 min. Animal movement was recorded using an overhead infrared-sensitive camera and analyzed using ANY-maze video-tracking software (version X; Stoelting Co., Wood Dale, IL, USA).

Visuospatial visual discrimination was evaluated using a water-maze-based task. Mice underwent 4 consecutive days of training, with 20 trials per day, to locate a submerged platform guided by the displayed visual stimulus. Correct-choice rates were recorded during training and under different spatial-frequency and contrast conditions.

Light-evoked avoidance behavior was evaluated in a two-compartment shuttle apparatus consisting of illuminated and dark chambers connected by an opening. Mice were habituated to the apparatus for 45 min on the preceding day. On the test day, each mouse was placed in the illuminated compartment and allowed up to 3 min to locate the dark compartment. Upon entering the dark compartment, a 15-min recording session was initiated under continuous illumination (447.5 nm, 5 mW/cm² at floor level). Transitions between compartments, latency to enter the dark compartment, and time spent in each compartment were recorded using an infrared-sensitive camera and analyzed with ANY-maze video-tracking software. Behavioral scoring was performed by investigators blinded to group allocation.

### Preprint status

This preprint reports the dataset available through November 16, 2026. Cohort expansion and additional validation experiments remain ongoing. Subsequent preprint versions will incorporate newly completed experiments, updated sample sizes, and finalized analyses.

## Results

### Characterization and extra-sheath transplantation of human retinal organoids (hROs)

The optic nerve was crushed 2 mm posterior to the globe, and day-30 hROs tissue was placed outside the optic nerve sheath at the injury site (Figure 1A, B). Before transplantation, donor tissue showed RAX and ISL1 immunoreactivity and spontaneous extracellular electrical activity recorded by MEA (Figure 1C), supporting retinal lineage differentiation and electrical excitability of the graft preparation.

**Figure 1.**
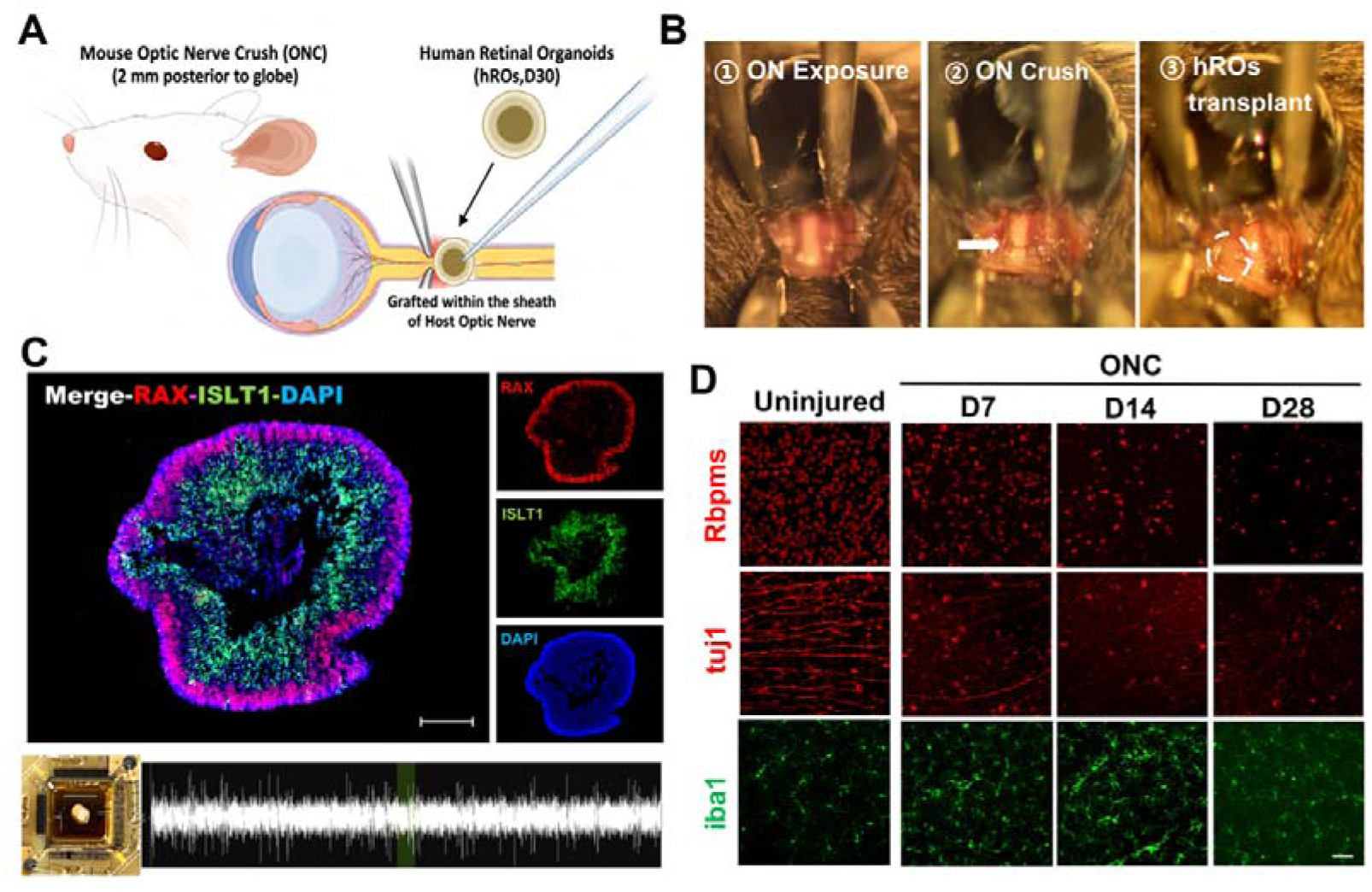
Generation of a mouse optic nerve crush model and hRO transplantation workflow. (A) Experimental schematic showing optic nerve contusion induced 2 mm posterior to the eyeball, with extra-sheath hROs implantation at the injury site. (B) Representative operative sequence showing optic nerve exposure, crush, and hROs transplantation. (C) Donor quality-control panel showing RAX and ISL1 immunoreactivity in day 30 hROs and a representative spontaneous microelectrode-array recording. Scale bars, 200 μm. (D) Time-series panel of retinal sections stained for RBPMS, TUJ1 and IBA1 after ONC (PSD7, PSD14, PSD28). Scale bars, 20 μm.

To characterize retinal degeneration after ONC, we examined retinal immunostaining at days 7, 14, and 28 after injury. Compared with control retinas, ONC retinas showed a progressive reduction in RBPMS-positive RGC somata, together with loss and disorganization of TUJ1-positive neuronal processes. IBA1 immunoreactivity also changed over time in its distribution and cellular morphology (Figure 1D). These findings were consistent with progressive inner retinal degeneration and an evolving retinal immune response after ONC.

### Transplantation partially preserves retinal structure and improves visual evoked responses

Fundus imaging revealed changes in optic disc appearance after ONC (Figure 2D). In ONC-only eyes, the optic disc appeared paler, its margins were less distinct, and peripapillary vessels were less conspicuous. Transplanted eyes showed better-preserved optic disc contours and more visible vascular branches. OCT demonstrated inner retinal thinning after ONC. GCL thickness did not differ significantly between the Trans and ONC groups at days 7 or 14, but was significantly greater in the Trans group at day 28 (Figure 2E). Thickness remained below control levels, indicating partial preservation of retinal structure.

**Figure 2.**
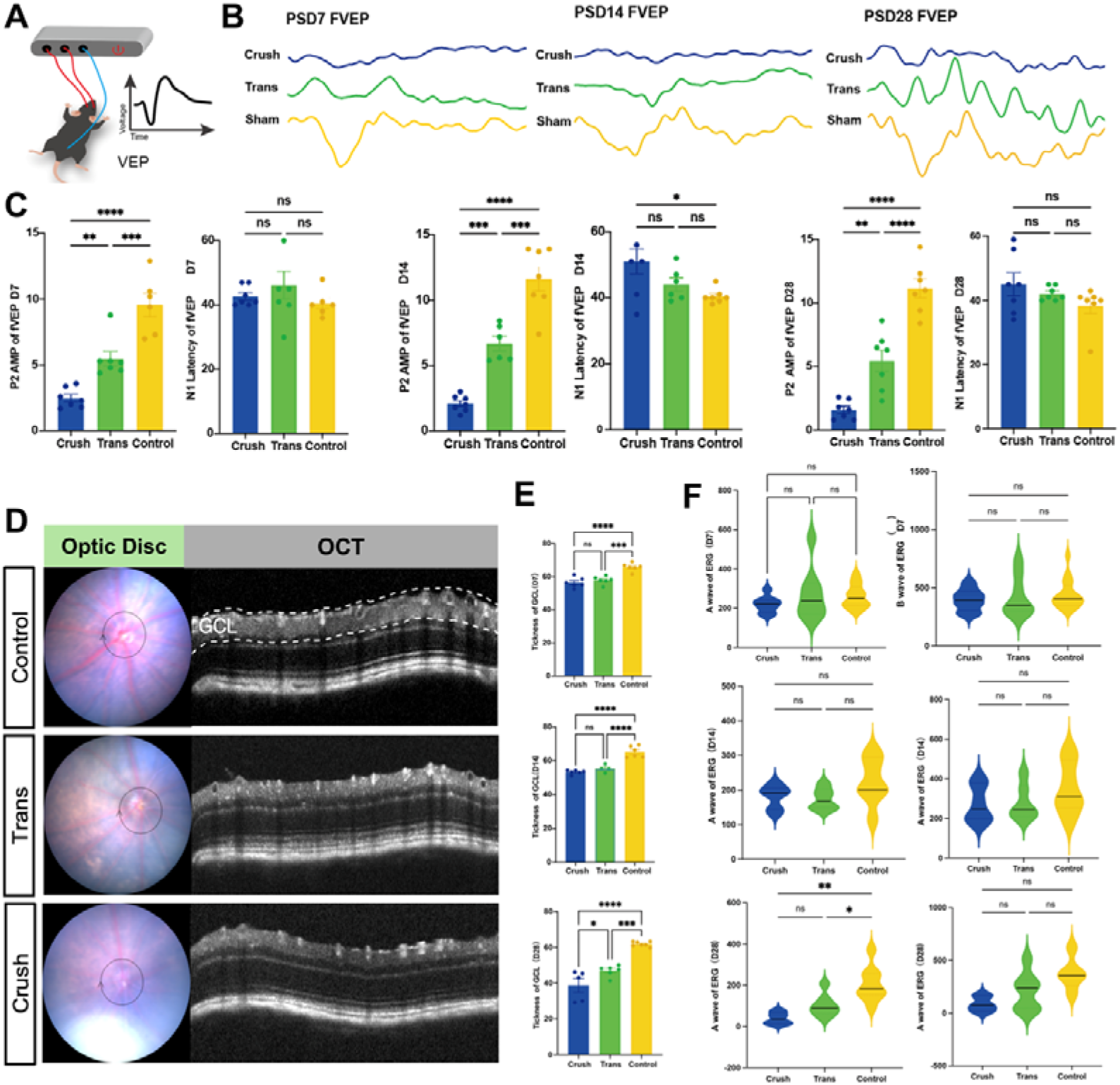
hROs transplantation partially preserves retinal structure and improves flash visual evoked potential responses after ONC. (A) Schematic of the flash VEP recording configuration. (B) Representative FVEP traces from crush, transplantation, and sham groups at post-surgical days 7, 14, and 28. (C) Quantification of P2 amplitude and N1 latency. (D) Representative optic-disc photographs and OCT B-scans identifying the ganglion cell layer. (E) GCL-thickness quantification at the same time points. (F) Full-field ERG a-and b-wave distributions. Data are presented as mean ± SD. n.s., not significant; *p < 0.05; **p < 0.005; ***p < 0.0005.

FVEP amplitude was reduced after ONC and was significantly higher in the Trans group than in the ONC group at days 7, 14, and 28, although it remained below control levels (Figure 2A-C). N1 latency generally did not differ significantly among groups, apart from a significant prolongation in the Crush group compared with the Control group at day 14 (p < 0.05), whereas the Trans group did not differ significantly from either group. Thus, transplantation was associated with increased evoked-response amplitude, while the latency results did not show a consistent treatment effect.

Full-field ERG a- and b-wave amplitudes did not differ significantly between the Trans and ONC groups at the measured time points (Figure 2F). These recordings did not detect an effect of transplantation on the fERG components assessed.

### Transplantation improves visual tracking and visually guided task performance

The OMR apparatus and grating stimuli are shown in Figure 3A. ONC mice exhibited reduced tracking responses across the tested spatial-frequency and contrast conditions (Figure 3B, C). Group comparisons showed impaired spatial vision after ONC and significantly better OMR performance in the Trans group than in the ONC group (Figure 3D), consistent with improved visually guided tracking after transplantation.

**Figure 3.**
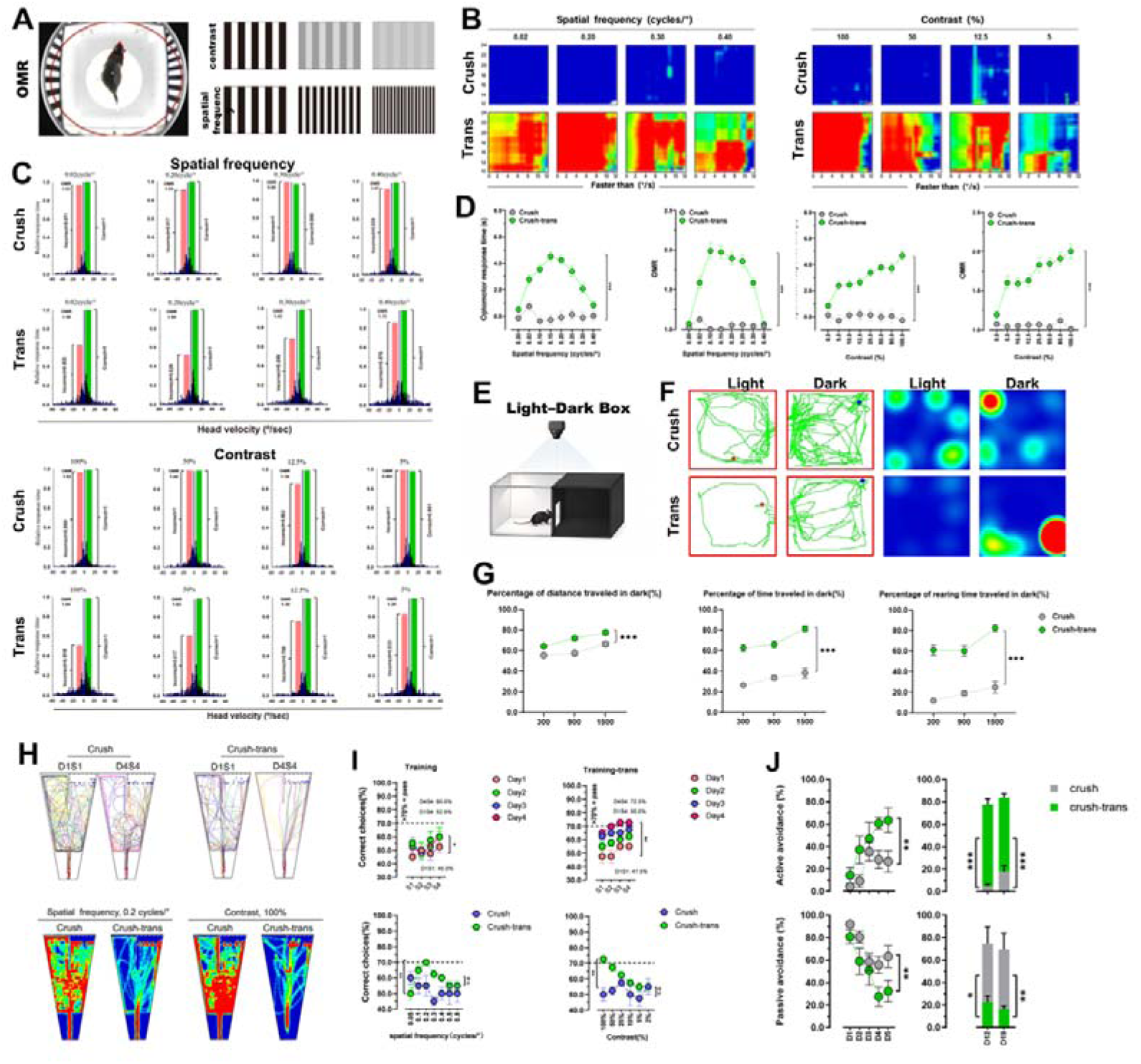
hROs transplantation improves selected visually evoked and light-context behavioral outcomes after ONC. (A) Optomotor-response apparatus and stimulus schematic. (B and C) Representative response distributions across contrast and spatial-frequency conditions. (D) OMR performance comparisons between crush and crush-transplantation groups. (E) Light-Dark Box apparatus. (F) Representative activity maps or trajectories at 1,500 lux. (G) Dark-compartment distance, time, and rearing as a percentage of total behavior at 300, 900, and 1,500 lux. (H and I) Visual-choice or training panels across spatial-frequency and contrast conditions. (J) Active and passive avoidance panels. Data are presented as mean ± SD. n.s., not significant; *p < 0.05; **p < 0.005; ***p < 0.0005.

We next assessed responses to graded illumination using the light-dark box (Figure 3E). Representative trajectories at 1,500 lux are shown in Figure 3F. ONC mice showed lower proportions of time, distance traveled, and rearing events in the dark compartment than controls across the tested illumination conditions. These measures were significantly increased in transplanted mice relative to ONC mice (Figure 3G). The findings indicate a change in behavioral responses to illumination after transplantation, although the assay does not isolate visual function from locomotor activity or anxiety-related behavior.

In the water-maze visual discrimination task, representative swimming trajectories and heat maps differed between ONC and transplanted mice (Figure 3H). Quantitative analysis showed lower correct-choice rates after ONC during training and under the tested spatial-frequency and contrast conditions. Transplanted mice achieved higher correct-choice rates than ONC mice (Figure 3I), indicating improved performance in a task requiring visual discrimination.

ONC also altered active and passive response proportions in the light-avoidance task. Compared with ONC mice, transplanted mice showed more active responses and fewer passive responses (Figure 3J). Together with the OMR and visual discrimination results, these observations were consistent with a behavioral benefit after transplantation. They do not, by themselves, establish recovery of learning or other cognitive functions.

### Transplantation preserves RGC somata and neuronal processes

RBPMS immunostaining showed progressive loss of RGC somata after ONC (Figure 4A). RBPMS-positive cell counts were similar in the Trans and ONC groups at day 7, but were higher in the Trans group at days 14 and 28 (Figure 4D). TUJ1 staining also indicated better preservation of neuronal processes after transplantation (Figure 4B). Total TUJ1-positive process length was greater in transplanted than in ONC-only retinas at each measured time point (Figure 4E).

**Figure 4.**
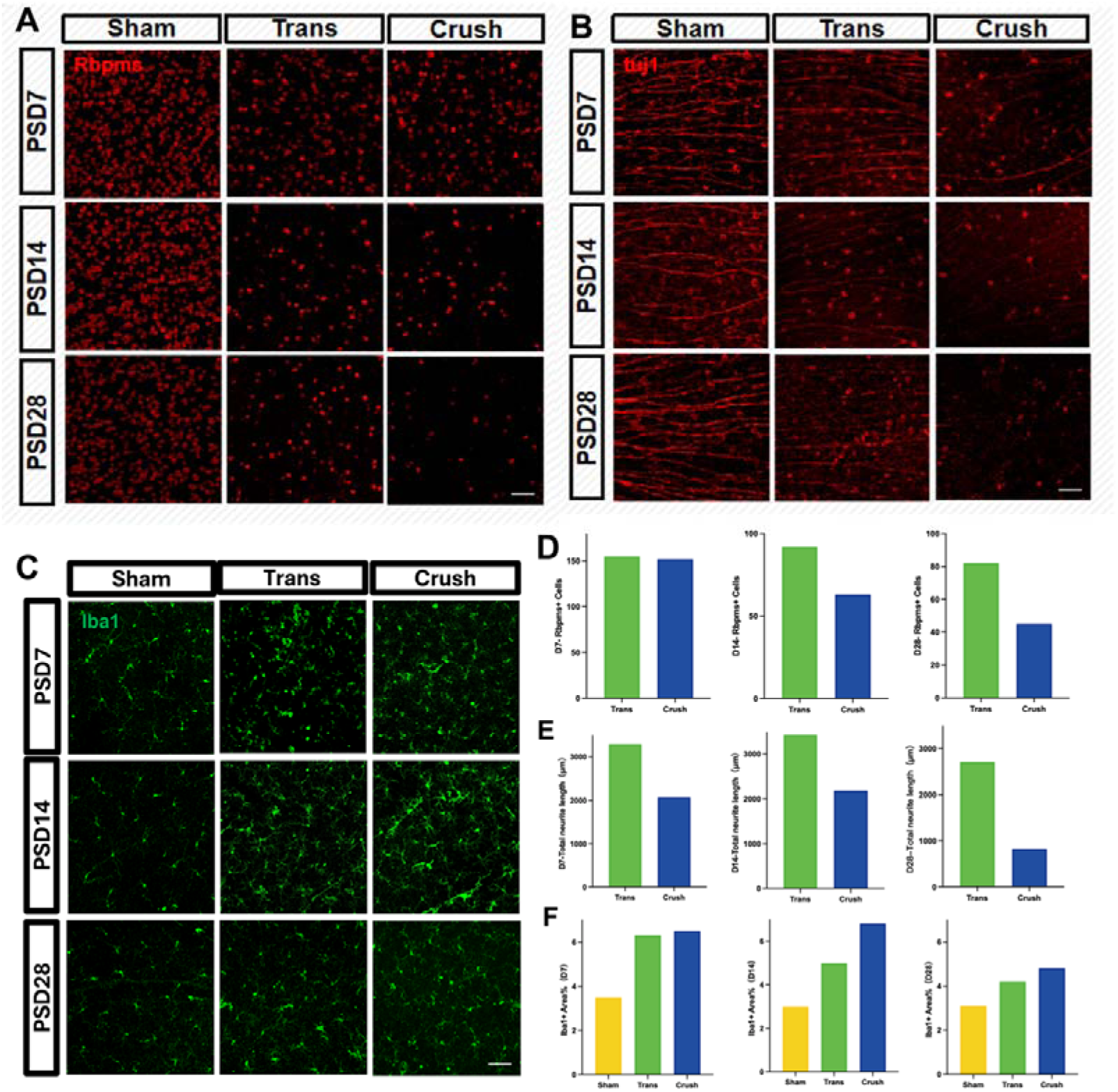
RGC-associated staining, neurite integrity, and microglial staining after hROs transplantation. (A) Representative longitudinal RBPMS staining in sham, transplantation, and crush groups. Scale bars, 20 μm. (B) Representative TUJ1 staining across the same groups and time points. Scale bars, 20 μm. (C) IBA1 staining after ONC and transplantation. (D) Displayed RBPMS-positive cell summary at days 7, 14, and 28. Scale bars, 20 μm. (E) Displayed total-neurite-length summary. (F) Displayed IBA1-positive-area summary. Sample size to be supplemented.

IBA1-positive area increased after ONC relative to sham surgery (Figure 4C). At later time points, transplanted retinas had a smaller IBA1-positive area than ONC-only retinas, although values remained above sham levels (Figure 4F). These findings associate transplantation with preservation of RGC somata and neuronal processes and with altered retinal IBA1 immunoreactivity after injury.

## Discussion

In the present study, extra-sheath transplantation of human retinal organoids (hROs) at the optic nerve crush (ONC) lesion site was associated with partial preservation of retinal structure and improvement in multiple measures of visual-pathway function. Before transplantation, donor hROs displayed retinal-lineage marker expression and spontaneous microelectrode-array activity (Figure 1C). Following ONC, hROs-treated mice showed a more preserved optic-disc appearance, delayed preservation of ganglion cell layer thickness, attenuation of retinal ganglion cell (RGC)-associated loss, enhanced FVEP responses, and improved performance in visually guided behavioral tasks (Figures 2–4). Together, these findings support hROs transplantation as a candidate multi-level reparative strategy after optic nerve injury. However, the current evidence supports partial functional recovery rather than complete reconstruction of the visual pathway.

The structural findings indicate that hROs transplantation primarily affected the injured inner retina. ONC induced progressive loss of RBPMS-positive RGC-associated somata, disruption of TUJ1-positive neuronal processes, and thinning of the ganglion cell layer, consistent with the delayed degeneration that follows RGC axonal injury^1, 18, 19^. hROs transplantation did not normalize retinal morphology, and GCL thickness did not differ from the ONC group at days 7 and 14. Nevertheless, the greater GCL thickness observed at day 28, together with improved preservation of RBPMS-positive cells and TUJ1-positive neurites, indicates attenuation of post-injury degeneration rather than complete reversal of injury-associated damage (Figures 2D, E and 4A, B, D, E).

The lower IBA1-positive area in hROs-treated retinas relative to ONC retinas at later time points further suggests that transplantation modified the injury-associated retinal microenvironment (Figure 4C, F). This finding is compatible with altered microglial activation or local inflammatory modulation. However, the present data do not define microglial phenotype or establish a causal relationship between IBA1 changes and functional recovery. Similarly, the current findings do not directly demonstrate inhibition of neuronal apoptosis and should not be interpreted as proof of a specific anti-apoptotic mechanism.

Electrophysiological findings further localize the treatment-associated benefit to the inner retinal and post-retinal visual pathway. hROs-treated mice showed higher FVEP P2 amplitudes than ONC mice across the measured time points, indicating improved visually evoked cortical responsiveness (Figure 2B, C). In contrast, full-field ERG a-wave and b-wave responses did not show a transplant-specific recovery relative to the ONC group (Figure 2F). This pattern argues against broad restoration of outer-retinal photoreceptor function and instead supports improved function within the RGC-to-visual-cortex axis. RGC survival, axonal regrowth, axonal conduction, and cortical responses represent related but non-equivalent outcomes after optic nerve injury^3–6, 22^.

Retinal organoids provide a biologically plausible source of retinal neural cells and may confer benefit through trophic support, modulation of the local microenvironment, or preservation of residual host-pathway activity. Previous studies have shown that pluripotent-stem-cell-derived retinal organoids and RGC-like cells can survive, differentiate, and provide functional or neuroprotective effects in experimental models^25–29^. However, the present study cannot distinguish among host-RGC neuroprotection, inflammation modulation, enhancement of residual visual-pathway activity, and donor-derived relay formation. Direct evidence of donor survival, donor axonal extension, synaptic integration, and connectivity with central visual targets is required before concluding that hROs reconstruct an eye-to-brain visual circuit.

The behavioral results provide convergent evidence that hROs transplantation improved visually guided function after ONC. Treated mice performed better in OMR testing across the assessed spatial-frequency and contrast conditions, supporting improved spatial vision and visually guided tracking responses (Figure 3A–D)^23^. hROs transplantation also improved dark-compartment behavior in the Light–Dark Box across graded illumination conditions and was associated with better performance in the water-maze-based visual discrimination and light-avoidance tasks (Figure 3E–J). These findings suggest that the treatment effect extended beyond basic visual detection to visually guided sensorimotor behavior, light-responsive behavior, and visual learning. Nevertheless, Light– Dark Box performance can also be influenced by locomotion, anxiety-like behavior, and motivation; it therefore supports, but does not independently prove, restoration of visual function^24^.

Overall, the available data support the conclusion that extra-sheath hROs transplantation after ONC is associated with partial preservation of RGC-associated structure, improved visual-pathway responsiveness, and amelioration of visually guided behavioral deficits. The results provide preliminary support for hROs transplantation as a potential experimental strategy for optic nerve injury. Expanded cohorts, independent replication, donor-specific tracing, synaptic validation, and rigorous analysis of graft-host interactions will be necessary to define the durability, mechanism, and translational relevance of this approach^30^.

## Limitations

The existing data support treatment-associated structural and functional benefit but do not independently establish donor-to-host synaptic integration or a regenerated eye-to-brain circuit. These limitations should be resolved before the work is submitted as a mechanistic cell-therapy study.

## Acknowledgments

This study was supported by funding from the National Key Research and Development Program of China (2024YFA1108700), and the Chongqing Science and Health Joint Major Project (Grant No. 2026DBXM007).

## Author contributions

Y.L., X.C. and H.X. designed the experiments and wrote the manuscript. Y.L. operated most of the experiments, wrote the main manuscript text, and integrated the figures. X.C. and H.Q. assisted with some of the experiments. Manuscripts and Figures were further verified by H.Q., J.K. and K.Y.. H.X. revised the manuscript and provided general guidance. All authors have read and approved the article.

## Data availability

Data will be made available on request.

## Ethics approval

This study was approved by the Institutional Animal Care and Use Committee of Army Medical University, Chongqing, China. All animal trials were approved by the Army Medical University Institutional Review Board, and all trials met guidelines.

